# Mitochondrial uncoupling protein 2 (UCP2), but not UCP3, is sensitive to oxygen concentration in cells

**DOI:** 10.1101/2020.07.01.181768

**Authors:** Karolina E. Hilse, Anne Rupprecht, Kristopher Ford, Olena Andrukhova, Reinhold Erben, Elena E. Pohl

## Abstract

One of the important hallmarks of cardiovascular disease is mitochondrial dysfunction, which results in abnormal energy metabolism and increased ROS production in cardiomyocytes. Members of the mitochondrial uncoupling protein family, UCP2 and UCP3, are thought to be beneficial by reducing ROS due to mild uncoupling. More recent hypotheses suggest the involvement of both proteins in cell metabolism by the transport of yet unknown substrates. The protein expression pattern under physiological and pathological conditions is an important clue for the evaluation of UCP2/UCP3 function, however, there is still no consensus about it. Previously, we demonstrated that only UCP3 is present in the adult murine heart under physiological conditions and correlated it with the predominant use of fatty acids for oxidation. In contrast, UCP2 was found only in very young (stem cell – like) cardiomyocytes, that rely mostly on glycolysis. Here, we employed three different models (ex vivo heart ischemia-reperfusion model, myocardial infarction model, and embryonic stem cell differentiation into cardiomyocytes under hypoxic conditions) to evaluate the abundance of both proteins under ischemia and hypoxia conditions. We found that (i) oxygen shortage or bursts did not influence UCP3 levels in the heart and ii) UCP2 was not present in healthy, ischemic, or re-perfused hearts. However, (iii) UCP2 was sensitive to the oxygen concentration in stem cells, in which UCP2 is normally expressed. These results further support the idea, that two highly homologous proteins – UCP2 and UCP3 – are abundant in different cells and tissues, and differently regulated under physiological and pathological conditions.

## 1. Introduction

Heart diseases continue to be a major cause of mortality worldwide [1]. Impaired energy metabolism and increased cellular and mitochondrial reactive oxygen species (ROS) are hallmarks that illustrate the critical role of mitochondria in the pathogenesis of heart disease [2]. Increasing evidence suggests that in addition to the generation of ATP and regulation of mitochondrial calcium and ROS, mitochondria are involved in dysfunctional cardiomyocyte metabolism during myocardial infarction and ischemia [3, 4].

Members of the uncoupling protein (UCP) subfamily – UCP2 and UCP3 – have been first proposed to modulate mitochondrial ROS production [5] based on their ability to transport protons. Meanwhile, it is becoming increasingly obvious that both proteins may be involved in cell metabolism by transporting other substrates [6-8]. The protein expression pattern is an important clue for the evaluation of UCP2 and UCP3 physiological and transport functions. However, no consensus about this issue exists in the community, mainly due to the low specificity of antibodies and the translational and posttranslational regulation of these proteins [9], that make the gene expression analysis less helpful [10]. Additionally, different physiological conditions and animal models yield contradictory results regarding the expression and function of UCP2 and UCP3.

UCP3, which was originally discovered and intensively studied in skeletal muscle [11], is found at even higher levels in cardiac muscle [12], making heart an important organ for UCP3 function analysis. Several groups have shown that a decrease or loss of UCP3 in rodent knockout models diminished contractile heart function and increased ROS production and tissue damage following ischemia, while its upregulation during ischemia improved the recovery of cardiomyocytes and increased cardiac efficiency [13-15]. In contrast, Nabben et al. failed to identify a protective role for UCP3 under lipid challenging conditions but showed that a lack of UCP3 led to a higher death rate in mice due to sudden cardiac arrhythmias [16]. Moreover, there is no consensus concerning the UCP3 expression pattern during heart failure. Young et al. and Razeghi et al. reported significant UCP3 mRNA downregulation in rodent and human failing heart [17-19]. Other groups detected UCP3 decrease under hypoxia in rat heart [20, 21]. In contrast, a positive correlation between protein abundance and heart ischemia was shown in other species [22-29].

Similar to UCP3, UCP2 expression in the healthy and failing heart is very controversially discussed [30, 31] that hindered the evaluation of its involvement in the pathogenesis of heart diseases. Both decreased [17, 18, 20] and increased [25, 26, 32-34] UCP2 abundance has been shown in different models of heart failure.

Unfortunately, most of these studies have in common that no specificity tests for the used antibodies were shown. Our group previously demonstrated that UCP2 is not present in the adult murine heart under physiological conditions [35, 36]. However, we detected UCP2 in very young (5-7 days) mouse hearts [36, 37], which is consistent with the concept that UCP2 is highly abundant in rapidly proliferating cells relying on glycolysis. In the hearts of young mice, UCP2 can be regarded as a marker of a cell “stemness” [37].

The goals of this study were to evaluate (i) the protein expression patterns of UCP2 and UCP3 under heart ischemia conditions and (ii) the impact of oxygen and ROS on the expression of both proteins at a cellular level. To this end, we used three different models: (1) *ex vivo* heart ischemia-reperfusion; (2) myocardial infarction model; (3) embryonic stem cell differentiation into cardiomyocytes under hypoxic conditions. We used custom-made antibodies against UCP2 and UCP3, which were evaluated in our previous works [35, 36].

## 2. Materials and Methods

### Animals

Three-month-old C57BL/6 wild-type (wt) mice were used in this study. Mice were kept under standardized laboratory conditions (12:12 h light-dark cycle, room temperature (24°C), food and water ad libitum). All animal procedures were approved by the Ethical Committee of the University of Veterinary Medicine, Vienna, and the Austrian Federal Ministry of Science and Research (permit number BMWF-68.205/0153-WF/V/3b/2014).

### Ex vivo heart ischemia-reperfusion model (Langendorff perfusion)

Following euthanasia by decapitation or cervical dislocation, the hearts of the mice were removed, and the coronary arteries perfused via the aorta at a constant pressure of 9.8 kPa at 37°C. Five different perfusion protocols were used to accommodate the various treatment conditions during the ischemia-reperfusion: 1) control – perfusion with Tyrode’s solution (116 mM NaCl, 20 mM NaHCO_3_, 0.4 mM Na_2_HPO_4_, 1 mM MgSO_4_, 5 mM KC1,11 mM D-glucose, and 1.8 mM CaCl_2_ [pH 7.4], gassed with 95% O_2_ and 5% CO_2_) for 5 min; 2) apocynin control – perfusion with Tyrode’s solution for 5 min followed by perfusion with Tyrode’s solution containing 100 μM apocynin [38] for 60 min; 3) ischemia control – perfusion with Tyrode’s solution for 5 min followed by 60 min global ischemia; 4) reperfusion – perfusion with Tyrode’s solution for 5 min followed by 60 min global ischemia and then reperfusion with Tyrode’s solution containing 100 μM apocynin for 60 min; 5) reperfusion control – perfusion as in (4) without apocynin. After the perfusion, the left ventricles were excised, snap-frozen in liquid nitrogen, and stored at −80°C.

### Myocardial infarction model

Acute myocardial infarction (AMI) was induced by permanent ligation of the left descending coronary artery (LDCA). In brief, mice were anaesthetised by intraperitoneal injection of ketamine/medetomidine (100/0.25 mg/kg) and placed under controlled ventilation with indoor air. Left lateral thoracotomy was performed at the fourth intercostal region (ICR), and the pericardium was removed to provide access to the left descending coronary artery. Ligation was performed 1 to 2 mm below the tip of the left atrial appendage. After the procedure, mice were treated for four days with Baytril^®^ and buprenorphine to alleviate pain and improve recovery. Mice were weighed weekly and monitored daily for excessive weight loss or abnormal behaviour. The sham operation was performed as above with the omission of coronary artery ligation. All mice were killed 4 weeks *post-surgery*.

### Western blot analysis

Heart samples were homogenised with a mixer mill in RIPA buffer supplemented with a protease inhibitor cocktail. After sonication and centrifugation, the lysate supernatant was collected. The total protein concentration was determined with the Pierce BCA Protein Assay Kit (Thermo Fischer Scientific). Total protein (20 μg) was separated on 15% SDS-PAGE gels and transferred to membranes (nitrocellulose membrane for UCP2 and PVDF membrane for UCP3). The membranes were incubated with antibodies against UCP3 and UCP2 (evaluated in [12] and [36] respectively), CII (SDHA, succinate dehydrogenase complex subunit A, Abeam, abl47l5, dilution 1:7500), GAPDH (glyceraldehyde-3-phosphate-dehydrogenase, Sigma Aldrich, G8795, dilution 1:7500), CV (ATP-Synthase, β-subunit, Invitrogen A-2135, dilution 1:5000), α-actin (Abcam, ab88226, dilution 1:3000), [3-actin (Sigma Aldrich, A5441, dilution 1:10000), Oct 3/4 (Santa Cruz, sc-9081, dilution 1:1500), and VDAC (voltage-dependent anion channel, Abeam, abl4734, dilution 1:5000). Detection was performed with the ChemiDoc-It 600 Imaging System (UVP, UK) using horseradish peroxidase-linked anti-rabbit and anti-mouse secondary antibodies (GE Healthcare, Austria) and ECL western blotting reagent (Bio-Rad, Austria). Western blot results were quantified with Launch Vision Works LS software (UVP, UK). The relative amounts of UCP2 and UCP3 were calculated as a ratio between the intensity of the protein band of the sample to that of the loaded mouse recombinant protein UCP2 or UCP3 [12]. The heart standard was prepared from pooled hearts of 2-5 months old C57BL/6 wt mice (n = 15) [37]. 20 mg of the heart standard was loaded per lane. The use of heart standard or recombinant proteins allowed the direct comparison of different western blots.

### Differentiation of mouse embryonic stem cells (mESC) into cardiomyocytes

mESC (clone D3, ATCC) were differentiated into cardiomyocytes according to the published protocol [39]. Stem cells were routinely maintained in Dulbecco’s modified Eagle’s medium (high glucose without sodium pyruvate; Thermo Fisher Scientific) supplemented with 15% fetal calf serum (Sigma-Aldrich), 2 mM L-glutamine (Thermo Fisher Scientific), 1% non-essential amino acids (Thermo Fisher Scientific), 0.1 mM [β-mercaptoethanol (Sigma-Aldrich), and antibiotics (50 U/mL penicillin/streptomycin; Thermo Fisher Scientific). To avoid stem cell differentiation and maintain pluripotency, the medium was supplemented with mLIF (10^6^ U/mL murine leukemia inhibitory factor; Millipore). mESC were split every 2 to 3 days. Cardiomyocyte differentiation was initiated by incubation of the mESC in maintenance media without mLIF for two days using the hanging drop culture method. The formed embryo bodies (EB) were transferred to bacterial Petri dishes and cultivated for an additional two days. EBs were then placed into 6-well plates and cultured for a maximum of 15 days. The contraction of the differentiated cardiomyocytes was determined by light microscopy. Cells were maintained in a humidified atmosphere under normoxic (5% CO_2_, 21% atmospheric O_2_) or hypoxic (5% CO_2_, 5% O_2_) conditions at 37°C. Reperfusion experiments were performed by transferring the cells from hypoxic to normoxic conditions. Cells were re-oxygenated for 3 days. The O_2_ tension under hypoxic conditions was controlled by an oxygen sensor and continuous injection of appropriate amounts of nitrogen.

### Statistical analysis

Western blotting results are presented as the mean ± SEM. Data were analysed using the t-test or one-way ANOVA for more than two groups (Sigma Plot 12.5 software). Data were considered statistically significant with a p < 0.05.

## 3. Results

### 3.1 Evaluation of the UCP3 expression in the ischemic heart

To evaluate the impact of short- and long-term exposure to hypoxia on the expression of UCP3 in the mouse heart, we performed two different sets of experiments on three-month-old mice: (i) Langendorff perfusion and (ii) acute myocardial infarction (MI). UCP3 expression was not changed following hypoxia with only a tendency to increase after reperfusion (Fig. 1, A). The relative amount of UCP3 (r.u.) was calculated as a ratio between the intensity of the protein band of the sample to the heart standard (Suppl. Fig. S1), which was prepared from 15 pooled hearts of 2–5 months old C57BL/6 wt mice (Hilse, Rupprecht et al. 2018). The equal amount of heart standard was loaded on each western blot that allowed the comparison of different western blots after the normalization of UCP3 to the heart standard.

**Figure 1.**
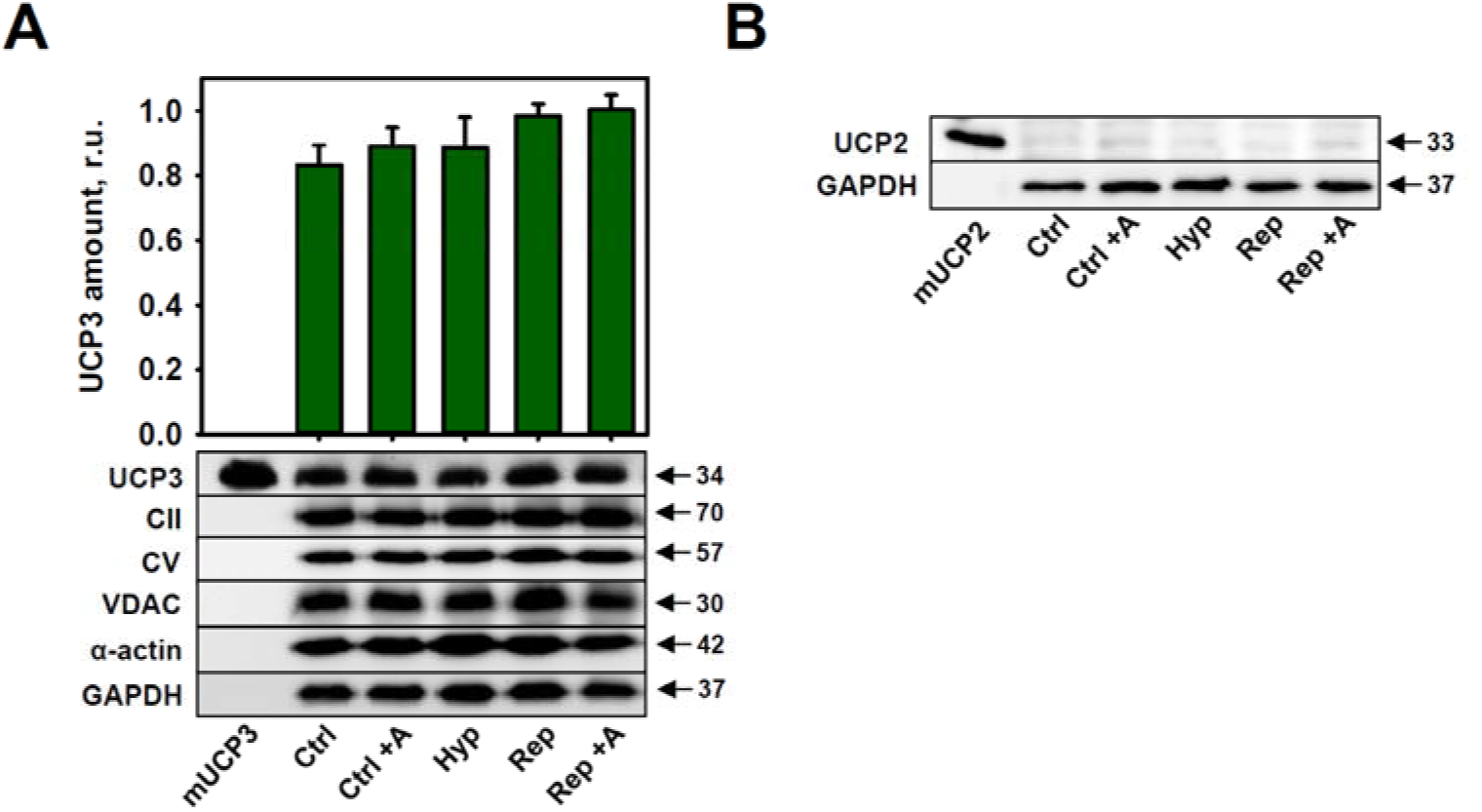
UCP2 and UCP3 expression in hypoxic heart model of Langendorff perfusion. (A) UCP3 expression in the murine heart under physiological conditions (Ctrl), during hypoxia (Hyp), reperfusion (Rep), reperfusion with apocynin (Rep+A). The relative amount (r. u.) of UCP3 is presented as a ratio between the UCP3 band intensity of a sample to heart standard (s. Materials&Methods and Suppl. Fig. S1). 20 μg of total protein from each heart sample were loaded per lane. 5 ng of the recombinant mUCP3 were loaded as a positive control (first lane of western blot). GAPDH and α-actin were used as loading controls. Data are presented as the mean ± SEM (n = 6 independent experiments). Numbers correspond to the molecular weight of investigated proteins. (B) UCP2 expression in the murine heart under physiological conditions (Ctrl), during hypoxia (Hyp), reperfusion (Rep), reperfusion with apocynin (Rep+A). 20 μg of total protein from each heart sample were loaded per lane. 2.5 ng of the recombinant mUCP2 were loaded as a positive control (first lane of western blot). GAPDH was used as loading control. A representative western blot from 6 independent experiments is shown.

To test the putative correlation between UCP3 expression and ROS production, we added a well-known inhibitor of NADPH oxidase and ROS production, apocynin (+A), to the perfusion buffer. Apocynin treatment during reperfusion did not affect the amount of UCP3 in the adult mouse heart (Fig. 1, A). We used different mitochondrial markers (CII, CV and VDAC [40]) to ensure that the mitochondria amount didn’t change under investigated conditions. UCP2 was neither present in the adult mouse heart under physiological conditions nor induced by short-term hypoxia or reperfusion (Fig. 1, B).

To examine the impact of long-term hypoxia on UCP3 levels in the mouse heart, we compared UCP3 expression levels in the left ventricle (LV; ischemic part of the heart) and right ventricle and septum (RV + S; normal perfused areas of the heart) of MI- and sham-operated animals, 4 weeks post-MI (Fig. 2, A and B). Western blotting showed high variability in UCP3 expression in both the left and right ventricles and septa across the MI mouse group (Fig. 2, A). UCP3 expression levels, normalized to those of the recombinant UCP3, revealed only a slight tendency to increase after MI (Fig. 2, B). Based on the high variance and lack of a statistically significant difference in UCP3 expression in the MI animals compared to the sham-operated controls, we concluded that oxygen stress does not appear to be the molecular mechanism that can cause the induction of UCP3 expression in cardiomyocytes.

**Figure 2.**
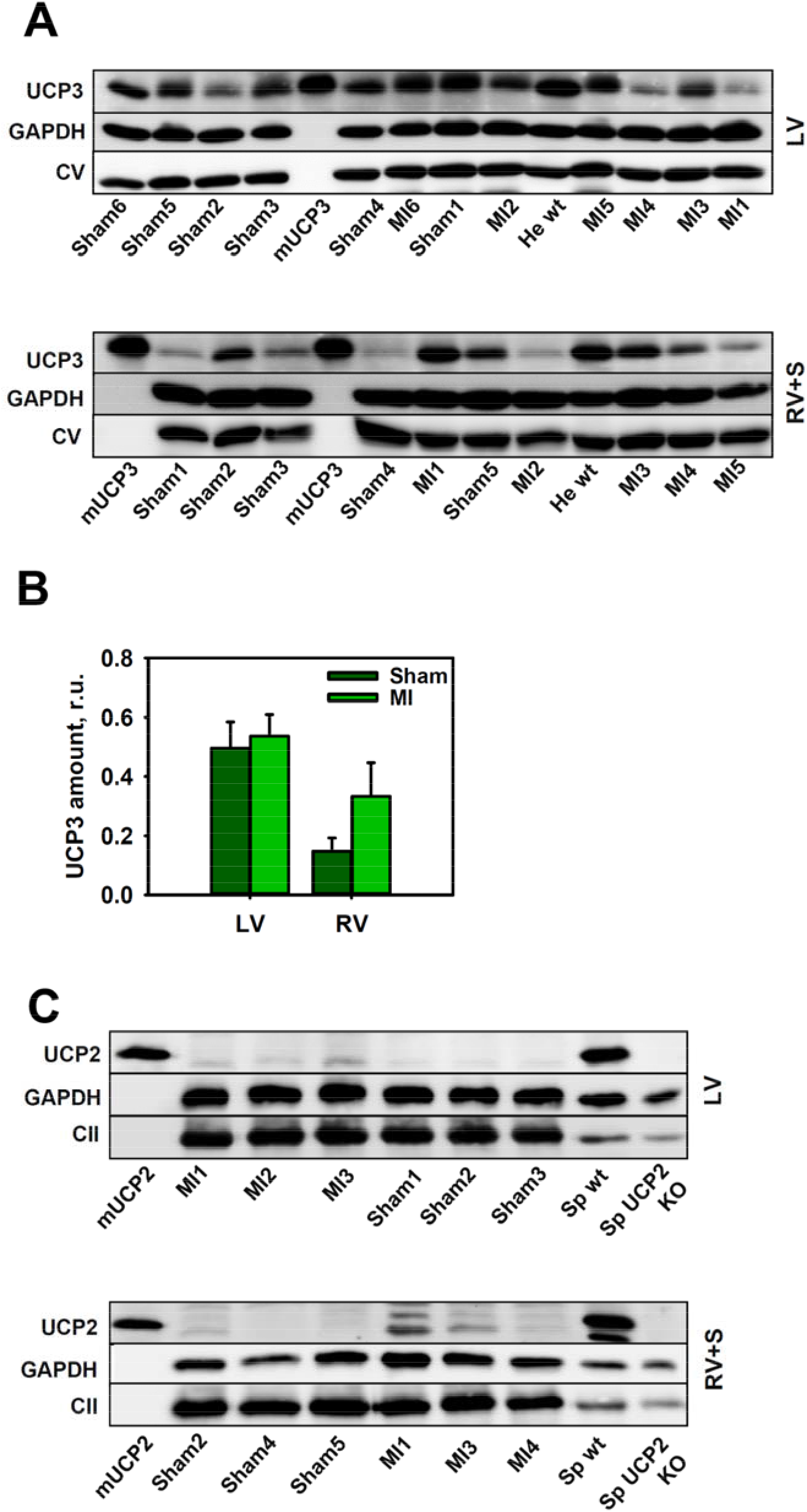
UCP2 and UCP3 expression after myocardial infarction in mice. (A) Representative western blots for UCP3 expression in the left ventricle (LV, upper blot) or right ventricle and septum (RV + S, bottom blot) after myocardial infarction (MI). Mice were divided into experimental (MI in LV and RV + S) and in control (sham in LV and RV + S) groups (n = 5 to 6 per group). 20 μg of total protein from investigated tissues were loaded per lane. mUCP3 (5 ng) and whole heart lysate from one mouse (He wt) were loaded as positive controls for UCP3. Antibody against CV was used as a mitochondrial marker, and GAPDH – as a protein loading control. (B) Quantification of UCP3 expression in the LV and RV+S from experimental and control mice. The relative UCP3 protein levels were calculated as a ratio of the intensity of the UCP3 band of the sample to that of the mUCP3. Data are presented as the mean ± SEM (n = 5 to 6 per group). (C) Representative western blots for UCP2 expression in the LV (upper blot) and RV+S (lower blot) after MI. 20 μg of total protein from investigated tissues were loaded per lane. mUCP2 (2.5 ng), whole heart lysate (He wt), or spleen lysate (Sp wt) were used as positive controls. Spleen lysate from UCP2KO mice served as a negative control. Antibody against CII was used as mitochondrial marker, and GAPDH was used as a protein loading control.

To test the hypothesis that UCP2 may serve as a marker appearing during oxygen deprivation in the adult heart, we performed western blotting on LV and RV + S samples from MI- and sham-operated animals. UCP2 was not expressed in the left ventricle of MI- or sham-operated animals. A slight increase in UCP2 expression was detectable in the right ventricle and septum after long-term hypoxia (Fig. 2, C). However, similar to UCP3 expression, we observed a high variance between animals, which might be also explained by inflammatory processes, e.g. the invasion of different numbers of immune cells [36, 41] or by adaptation of cellular metabolism [6].

### 3.2 Influence of oxygen levels and the induction of cardiomyocyte differentiation on UCP expression in stem cells

Recently, we demonstrated that UCP3 is not expressed in mESC under normal conditions as well as at any stage of mESC differentiation into cardiomyocytes [37, 42]. To exclude the possibility that UCP3 expression may be induced in the cardiomyocyte differentiation model after exposure to hypoxia, we incubated cells from day 5 to day 10 in hypoxic conditions.

Neither hypoxia nor re-oxygenation led to UCP3 expression, and such treatments did not change UCP2 abundance in the mESC-derived cardiomyocytes on day 10 (Fig. 3, A). Interestingly, UCP2 expression was significantly increased in stem cells, which are a natural habitat for UCP2 [42], under hypoxic conditions and then declined following re-oxygenation (Fig. 3, B).

**Figure 3.**
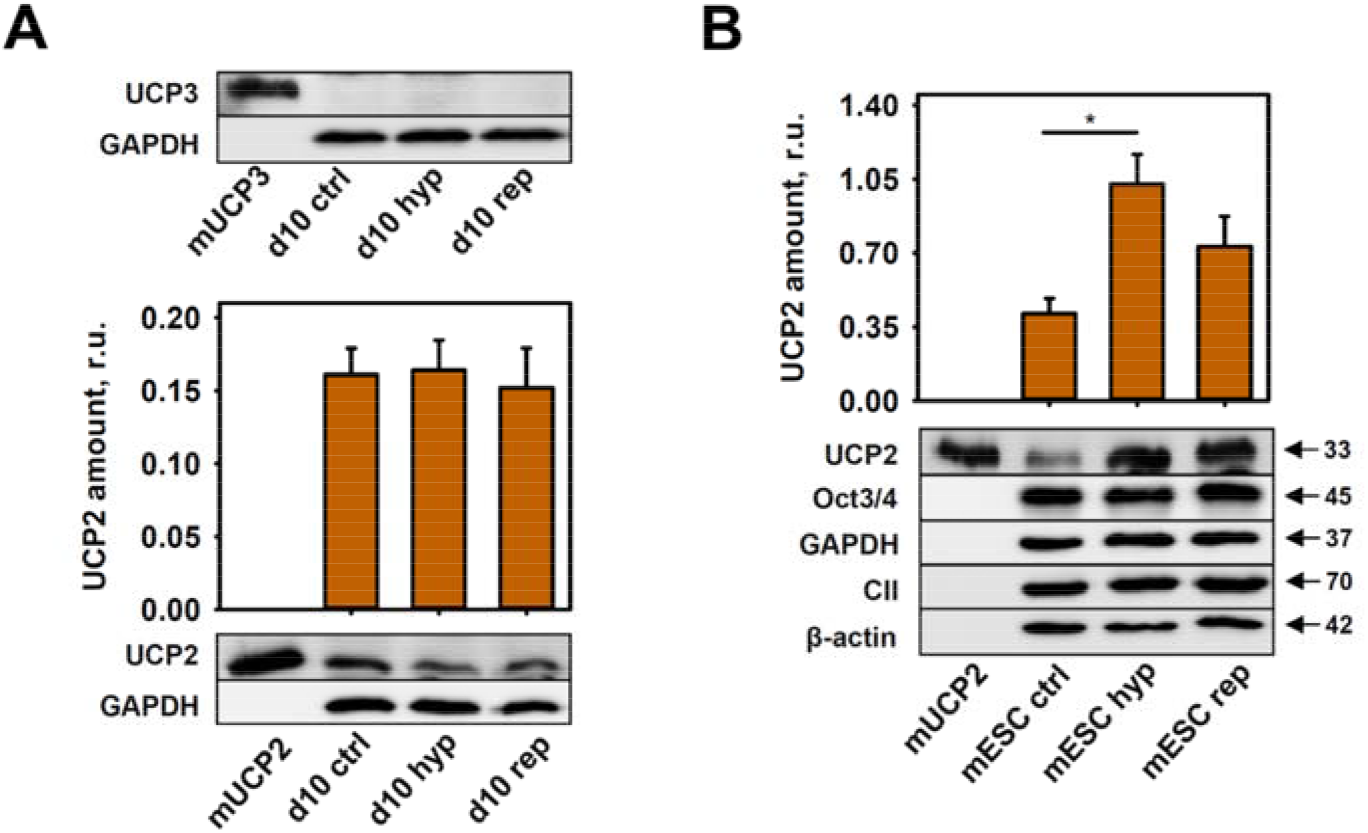
UCP2 and UCP3 expression in differentiated cardiomyocytes and mesenchymal mESC during hypoxia and reperfusion. **(A)** Representative western blotting analysis of UCP3 and UCP2 protein levels in mESC-derived cardiomyocytes (day 10) under standard conditions (ctrl) or during hypoxia (hyp) or reperfusion (rep). (B) Representative western blots and quantification of UCP2 expression in mESC under standard conditions (ctrl) or during hypoxia (hyp) or reperfusion (rep). 20 μg of total protein from cell lysates were loaded per lane. The relative UCP2 amount (r. u.) is a ratio of the UCP2 band intensity of the sample to that of recombinant UCP2 (2.5 ng). Data are presented as the mean ± SEM (n = 3-8 cell passages). An antibody against OCT3/4 was used as a stem cell marker; CII, β-actin, and GAPDH were used as controls for mitochondria amounts and protein loading.

## 4. Discussion

Understanding the complex role of the mitochondrial uncoupling proteins (UCP2 and UCP3) in the genesis and progression of cardiovascular disease (e.g., ischemia or heart failure) could provide novel targets for therapeutic intervention. Despite over two decades of intensive research, there is no consensus on the roles of UCP2 and UCP3 in the heart. Nevertheless, it is now becoming clear that no reasonable conclusions can be made based solely on UCP2 and UCP3 gene expression, because both proteins are subjected to strong translational regulation and have a short half-life [6, 30, 36, 43]. In addition, the expression of UCP2 and UCP3 is confined to tissues or cells with different type of metabolism, glycolysis and fatty acid oxidation respectively [10].

Since the discovery of UCP3, the majority of studies have focused on evaluating its putative function in the regulation of ROS production. In particular, UCP3 was discussed as a mild uncoupler, preventing damage to myocardial tissue in cardiovascular disease [24]. Indeed, UCP3 has proton-transporting activity, similar to UCP1 and UCP2 [44, 45]. Previous observations made with MI model in UCP3KO mice demonstrated larger infarcts, higher ROS production, and impaired myocardial energetics [14, 29]. It has been shown that the hearts of UCP3KO animals are not protected by ischemic preconditioning. The restoration of uncoupling (e.g., pharmacologically with carbonyl cyanide-4-(trifluoromethoxy)phenylhydrazone (FCCP) or via UCP1 overexpression in heart) was claimed to protect against IR injury [46], which appeared to confirm the protective role of UCP3 on the heart through mild uncoupling [47]. Several groups reported UCP3 upregulation by oxidative stress [48-51]. Jiang et al. [48] explained the increased UCP3 in skeletal muscle during exercise by the increased ROS generation. Interestingly, due to prolong exercise skeletal muscles switch to the increased use of fatty acids that positively correlate with the expression of UCP3 [10].

Our results on UCP3 expression in short- and long-time models of oxygen shortage (Langendorff perfusion and myocardial infarction) are not in line with the previous observations. The high variance and lack of a statistically significant difference in UCP3 expression in the MI animals in our study suggest that oxygen stress and ROS generation is not the global mechanism for the induction of UCP3 expression. This conclusion contradicts studies reporting UCP3 upregulation by oxidative stress in heart, but is in line with the view that “cardiac muscle may remain reliant on fatty acids during sustained hypoxia” [52]. The direct correlation between the expression of UCP3 and FAO was demonstrated previously [37, 53, 54]. For accuracy, it should be mentioned that reactive species can increase the protonophoric function of UCPs in the presence of fatty acids without affecting their expression [55, 56].

Our present results revealed that, in contrast to UCP3, UCP2 levels are sensitive to changes in the oxygen levels in mESC (Fig. 3, B). Our laboratory and few other groups previously demonstrated that UCP2 protein, whose mRNA expression is ubiquitous [57, 58], is solely present in highly proliferating cells that have a glycolytic type of metabolism (e.g., cancer, stem cells, and activated immune cells) [36, 42, 59, 60]. Therefore, our results that showed an increase in the UCP2 levels in mESC during hypoxia are in line with the observations that stem cells are more glycolytic under oxygen deprivation conditions and remain undifferentiated longer with an increased proliferation capacity [61]. We recently showed that the bioenergetics profile of mESC and mESC-derived cardiomyocytes revealed a highly glycolytic type of metabolism [37], which plausibly explains the high levels of UCP2 in these cells and in hearts of young mice as well as its changing expression levels during differentiation and aging [37, 42]. Similarly, the presence of UCP2 in the ischemic heart, although only in parts of the right ventricle and septum, could be explained by the presence of glycolytic, rapidly proliferating cells. Activation of resident cardiac stem cells to proliferation and differentiation after ischemic injury contributes to the tissue regeneration [62-64]. Additionally, the high levels of ROS that appear during reperfusion attract pro-inflammatory neutrophils [65]. As a consequence, immune cells, in which UCP2 is usually highly expressed, infiltrate damaged tissues [36, 41]. Nevertheless, metabolic adaptation to proliferation and oxidative stress is a dynamic and flexible process to constantly adjust to the actual situation. Such adaptation could be ensured by fast UCP2 expression as has been previously shown for neuroblastoma cells [6].

In summary, in this study, we have revealed that UCP3 expression is not altered by myocardial infarction or reperfusion. UCP2 levels were sensitive to changes in oxygen levels only in cells where it is typically present (i.e., stem cells). The results of this study support the hypothesis that two highly homologous proteins – UCP2 and UCP3 – are abundant in different cells and tissues, and differently regulated under physiological and pathological conditions. The re-evaluation of the previous results with regard to these new insights would be an important step for uncovering the exact functions of UCP2 and UCP3 in cardiomyocytes and their roles in cardiac dysfunction.

## Author Contributions

conceptualization, E.E.P., O.A. and K.H.; methodology, K.H., K.F., A.R., R.E.; formal analysis, K.H. and K.F; investigation, K.H. and K.F; writing—original draft preparation, K.H.; writing—review and editing, E. E. P., K.H., A.R., R.E.; funding acquisition, E.E.P, O.A.

## Acknowledgments

We are grateful to Sarah Bardakji, Ute Zeitz (Institute of Physiology, Pathophysiology and Biophysics, University of Veterinary Medicine, Vienna, Austria) for their excellent technical assistance. We thank Svetlana Slavic (Institute of Physiology, Pathophysiology and Biophysics, University of Veterinary Medicine, Vienna, Austria) for help with mouse surgery.

