## Supplementary material for "Mitochondrial uncoupling protein 2 (UCP2), but not UCP3, is sensitive to oxygen concentration in cells": Suppl. Figure S1

**Supplementary Figure**
**Hilse et al. (2020)**

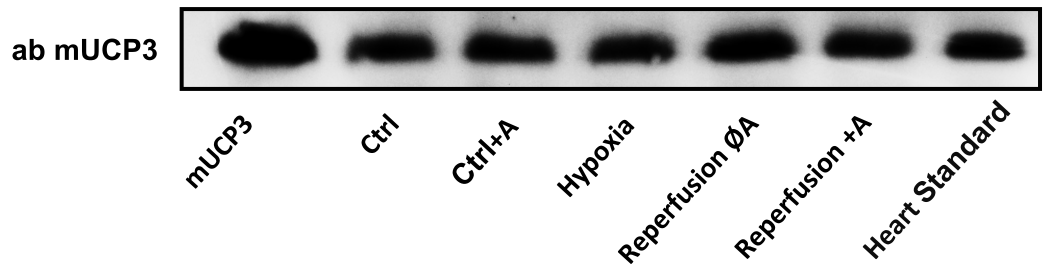


**Suppl. Figure S1.** UCP3 expression in hypoxic heart model of Langendorff perfusion. (A) UCP3 expression in the murine heart under physiological conditions (Ctrl), during hypoxia (Hyp), reperfusion (Rep), reperfusion with apocynin (Rep+A). Heart standard was prepared from pooled hearts of 2–5 months old C57BL/6 wt mice (n = 15) as described in (Hilse, et al. 2018, Frontiers Physiol.) and was used for the normalization of UCP3 amount (s. Fig. 1, A).
